# Production of Multilayer Cell Mass from Olfactory Stem Cell and Its Utilization Potential in Nervous Tissue Regeneration

**DOI:** 10.1101/102434

**Authors:** Olga N. Oztel, Adil Allahverdiyev, Aysegul Batioglu Karaaltin, Melahat Bağırova, Ercument Ovali

## Abstract

**Background:** Experimental studies performed with human olfactory nerve stem cells have shown that these cells can ameliorate nerve cell regeneration. Developing a method of repairing nerve damage solely using stem cells without the need of any supporting material is important.

**Methods:** A multilayer cell mass was obtained from olfactory tissue-derived mesenchymal stem cells with high viability and proliferation capability using a protocol devoid of scaffolds or any other artificial supporting material. First, human olfactory tissue-derived mesenchymal stem cells were isolated, cultured, and characterized. Next, consecutive passages were conducted to obtain multilayer cell growth. The resulting cell mass could be suitable for tissue engineering models as well as nerve cell or tissue regeneration studies in the future.

**Results:** Viability and adhesive properties of the resulting cell mass were examined and found to be suitable for use in nerve tissue regeneration.

**Conclusion:** It is suggested that an *in vitro*-produced olfactory stem cell mass can be applied to a very small damaged region and could have a high potential for microenvironment formation.

## 1. Introduction

Peripheral nerve damage can be caused by congenital abnormalities, tumor resection, and total transection. This may lead to neurological function loss that is difficult to restore (Akyildiz, 2002). Like other mammals, human neurogenesis develops through the olfactory bulb, creating neurons and interneurons at the subventricular region. The olfactory mucosa contains the epithelial layer (olfactory epithelium), lamina propria (olfactory glands), olfactory sensory axons, and specialized ensheathing cells. It is located in the dorsal and posterior part of the nasal cavity in mammals, medially covering the nasal septum and laterally covering the turbinate bones. It therefore provides a considerably large surface for sensory cilia that project from the dendrites of the sensory neurons. The multipotent stem cells that are located in the olfactory mucosa can be biopsied and isolated (Mackay-Sim, 2010; Curtis et al., 2009; Mackay-Sim, 2013). Human olfactory tissue-derived stem cells (OTD-SCs) have the ability to survive in a neurosphere culture. They are capable of differentiating into neuron, astrocyte, oligodendrocyte, and even non-neural cell types upon proper inductions (Curtis et al., 2009; Hidaka et al., 2013; Mackay-Sim, 2013; Murrell et al., 2008; Papanikolaou et al., 2008). These OTD-SCs were cultured in a neurosphere, differentiated, and used in the treatment of Parkinson’s disease. The formation of dopaminergic neurons was observed *in vitro*, and recovery was observed *in vivo* in rat (Hidaka et al., 2013; Mackay-Sim, 2013; Murrell et al., 2008; Papanikolaou et al., 2008). Previous studies have shown that stem cells can also be differentiated into cells similar to the Schwann cells that are responsible for myelination in the peripheral nervous system. These results indicate that these cells can be used in nerve regeneration by transplantation of autologous cells (Dong and Yi, 2010). All of these studies demonstrate the potential for OTD-SCs in nerve regeneration. There is a definite potential use for OTD-SCs in tissue engineering (Salehi et al, 2009; Huang et al., 2010; Leung et al. 2007; Ao et al., 2007; Wang et al., 2011; Khoo et al., 2011). However, there are a limited number of studies focused on the culture and *in vivo* studies of these cells. A number of problems arise when the cells are transferred *in vivo*. One of the main difficulties is that the cells in suspension cannot be properly localized to the damaged region. To overcome this difficulty, several methods have been applied for appropriate localization. The most common method is transferring cells by a Hamilton syringe. Although localization of the cells to the region of interest is easy with a steel needle, there is a risk of bursting the cells with a high injection pressure (Ajalloueyan et al., M, 2013; Martinez et al., 2009; Roussos et al., 2005). Multiple types of scaffolds have been used in tissue engineering as well as with OTD-SCs, but the preparation of the necessary materials and utilizing them for nerve regeneration is difficult and expensive (Ao et al. 2007; Ferrero-Gutierrez et al., 2013). Additionally, scaffolds take longer to biodegrade and can be toxic (Jayakumar et al., 2011; Yim et al., 2007; Haile et al., 2008).

This study is unique, as there are no other studies that have used multilayer cultures to produce viable olfactory tissue-derived mesenchymal stem cells that can be used in the regeneration of nerve tissue with no requirement of additional materials.

## 2. Methods

### 2.1. Isolation, culture, and cryopreservation of human OTD-SCs

Ethical approval was received from the Samatya Training and Research Hospital and performed in accordance with the Helsinki Declaration of the World Medical Association. Samples of the olfactory mucosa with a 0.3cm diameter were obtained from three male donors and transported to the Cell Culture and Tissue Engineering Laboratory of Yildiz Technical University’s Bioengineering Department in cold conditions. The tissue samples were washed twice with a DMEM/F12 solution in the center well. They were then incubated at 37ºC for 30 minutes in an 0,1% dispase II (m/v, Roche Diagnostics, Mannheim, Germany) solution in order to dissociate the tissue pieces. The tissue pieces were then transferred into a cell culture solution of DMEM/F12 and an antibiotic-antimycotic cocktail using sterile forceps. The pieces were further separated mechanically using insulin injector needles, whose tips were bent as 90°. These layers were then transferred into 6-well cell culture dishes. The culture dishes were covered with 12mm diameter cover slips and incubated in an DMEM/F12 medium that included 2 mM L-glutamine (Invitrogen, Netherlands), 1 % antibiotic– antimycotic solution (Life Technologies), and 10% fetal bovine serum (FBS) (Life Technologies). The medium was renewed every three days and the cellular adhesion was observed for 2 weeks.

At the end of the incubation period, the cover slips were removed and inverted using sterile forceps. They were placed in a new dish and the cells were led to proliferate. When 70% confluency was obtained on the cover slips, they were trypsinized and transferred into 25cm^2^ flasks. Cells were subcultured with Trypsin (0.05%)-ethylenediamine tetraacetic acid (EDTA, 1 mM) from Invitrogen.

### 2.2. Characterization of human OTD-SCs

#### 2.2.1. Immunostaining and Fluorescence-Activated Cell Sorting (FACS) Analysis

Flow cytometry analyses were performed using the BD FACSCalibur (BD Biosciences, San Jose, CA) following the third passage. After the cells were trypsinized, they were counted and homogenized in phosphate buffered saline (PBS). Cells were incubated with 10µl of fluorescein isothiocyanate (FITC) or phycoerythrin (PE)-conjugated monoclonal antibodies (BD Biosciences) (CD14-PE (cat no:557154), CD117-PE (cat no:561682), CD11b-PE (cat no:557321), CD13-PE (cat no:555394), CD44-PE (cat no: 555479), anti-human CD90 (cat no: 555596), anti-human CD116-PE (cat no:551373), Mouse IgG1-PE (cat no:550617), CD146-PE (cat no: 550315), CD73-PE (cat no: 550257), HLA-ABC-PE (catno555553), CD29-PE (catno: 561795) (BD Biosciences), Mouse IgG1 K- PE (cat no:555749), CD105-PE (cat no:560839), Mouse IgG1- FITC (cat no:555909), anti-HLA-DR-FITC) (cat no:556643) at RT in the dark for 45 minutes. After incubation, cells were centrifuged in a PBS solution containing 0,1% sodium azide for 5 min at 1780 rpm. They were then re-suspended with 400µl of cell washing buffer. Prepared cells were processed with the BD FACSCalibur flow cytometry device and the results were analyzed using the BD Cell Quest^TM^ software.

#### 2.2.2. Differentiation of OTD-SCs into Adipocytes

Isolated OTD-SCs were differentiated into adipocyte cells by using an 2ml STEMPRO^®^ Adipogenesis Differentiation Medium (Invitrogen). During differentiation, complete medium changes were made every 3 days. After 14 days, cultures were monitored for differentiation by using Oil Red O (Sigma) (0,5% Oil Red O solution in 60% isopropanol), washed with distilled water, and visualized.

### 2.3. Staining of OTD-SCs with Chloro-methylbenzamido dialkylcarbocyanine (CM-DiI)

CM-DiI (1 g/L) (Invitrogen) was prepared in dimethyl sulfoxide (DMSO). OTD-SCs with 70% confluency were washed with PBS two times and trypsinized. OTD-SCs suspension (500 μL) was prepared by adding OTD-SCs (5×10^6^) of the 3^rd^ passage to PBS. Five microliter CM-DiI was added into 500 μL OTD-SC suspension and incubated at 37°C for 4 min and 4°C for 15 min. The CM-DiI-labeled OTD-SCs were washed using PBS for two times, re-suspended in 50 μL PBS, and seeded into 25cm^2^ tissue culture flask. The growth of CM-DiI-labeled OTD-SCs were evaluated under a fluorescence microscope using ordinary light and light of a wavelength of 540-650 nm.

### 2.4. Formation of Multilayer Cell Pellet

In order to prepare the multilayer cell mass, CM-DiI-labeled OTD-SCs with 70% confluency were removed from the surface using a trypsin-EDTA solution. These trypsinized cells were placed in a new 25cm^2^ flask (the master culture). This step was repeated 8 times every other day, with the cells being accumulated in the same master culture flask. The resulting cell mass was then transferred into a sterile petri dish containing PBS and an antibiotic-antimycotic solution. The multilayer cell mass was cut into 1mm^3^ pieces using sterile forceps and transferred into a 6-well petri dish to detect the invasion ability of these cells. After covering the dish with 12mm^2^ cover slips, the cells, in a DMEM/F12 medium containing 10% FBS, were incubated at 37°C in a 5% CO_2_ incubator.

## 3. Results

### 3.1. Obtaining olfactory mucosa and isolation of stem cells

The three different donors were 35, 36, and 40 years old. There was no variation among the characteristics of the cell cultures obtained from the donors. Cell adhesion was observed in the olfactory tissue following primary culture in the second week. After 3-4 weeks of cultivation in DMEM/F12, a subculture of the cells was then performed (Fig. 1).

**Figure 1.**
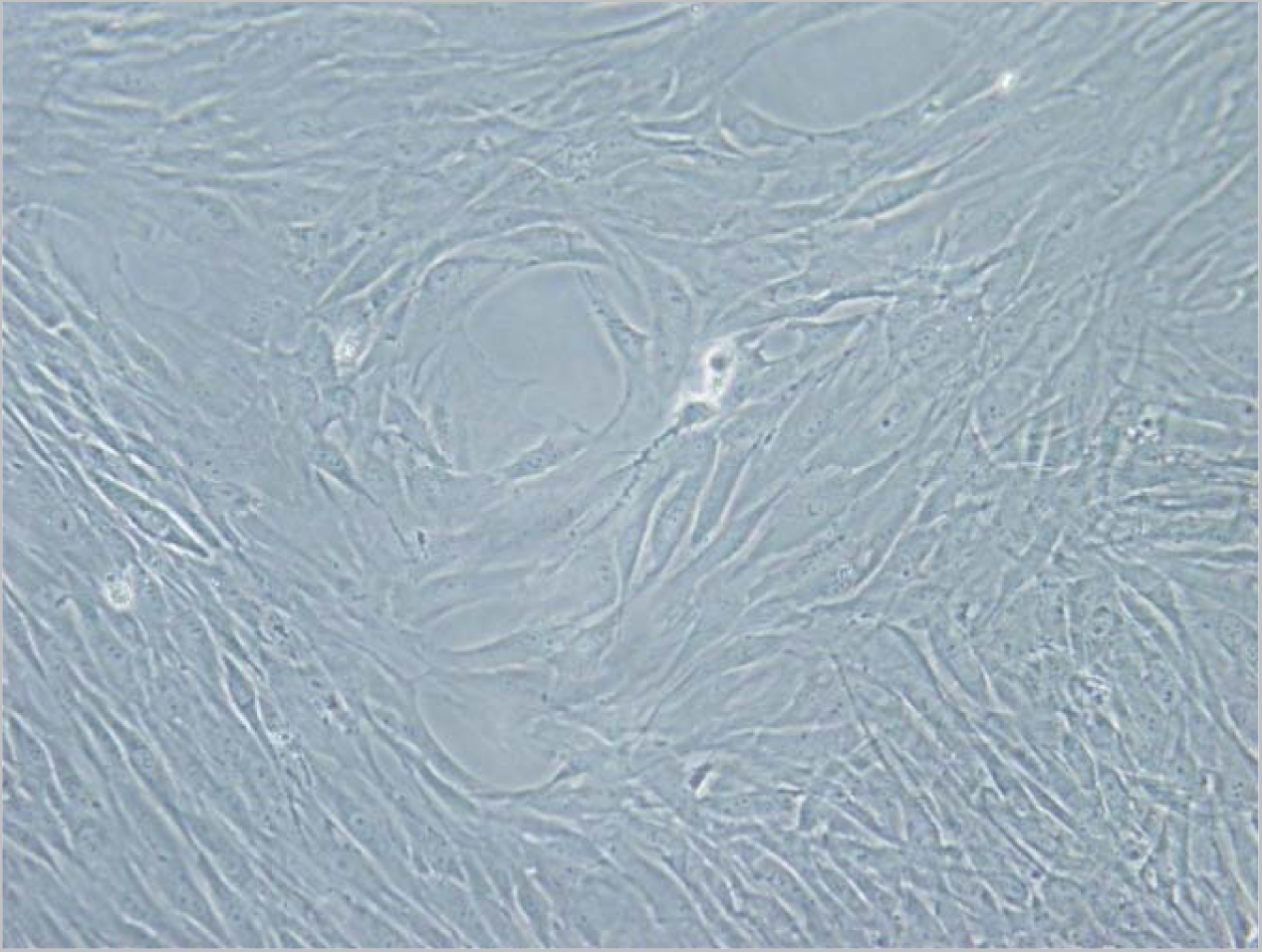
Cultured olfactory stem cells on the 21th day (20X)

### 3.2. Flow cytometric analysis of OTD-SCs

As mentioned in the methods section, the following MSC specific cell surface markers were analyzed by using either FITC- or PE-conjugated antibodies on the stem cells: CD14-PE, CD117-PE, CD11b-PE, CD13-PE, CD44-PE, anti-human CD90, anti-human CD116-PE, Mouse IgG1-PE, CD146-PE, CD73-PE, HLA-ABC-PE, CD29-PE, Mouse IgG1 K-PE, CD105-PE, Mouse IgG1-FITC, and anti-HLA-DR-FITC (Fig. 2). These findings were similar to the immunophenotypic MSC characteristics of OTD-SC reported before (Batioglu-Karaaltin et al., 2016).

**Figure 2.**
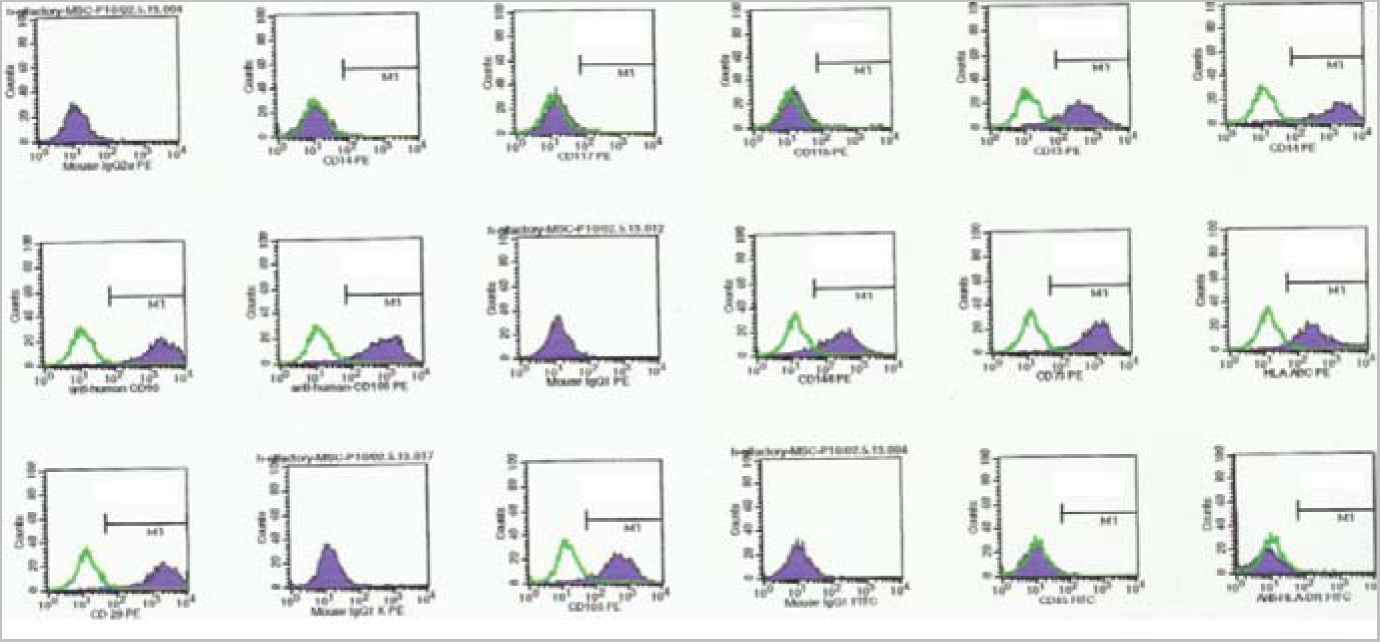
Immunophenotyping results of cultured olfactory mucosa derived stem cells. The figure shows the corresponding distribution of each CD marker anayzed (CD14-PE, CD117-PE, CD11b-PE, CD13-PE, CD44-PE, anti-human CD90, anti-human CD116-PE, Mouse IgG1-PE, CD146-PE, CD73-PE, HLA-ABC-PE, CD29-PE, Mouse IgG1 K-PE, CD105-PE, Mouse IgG1-FITC, anti-HLA-DR-FITC respectively).

### 3.3. Staining stem cells with CM-DiI

Successfully stained cells were analyzed using a phase-contrast microscope and UV light with the green filter according to the manufacturer’s protocol (Fig. 3).

**Figure 3.**
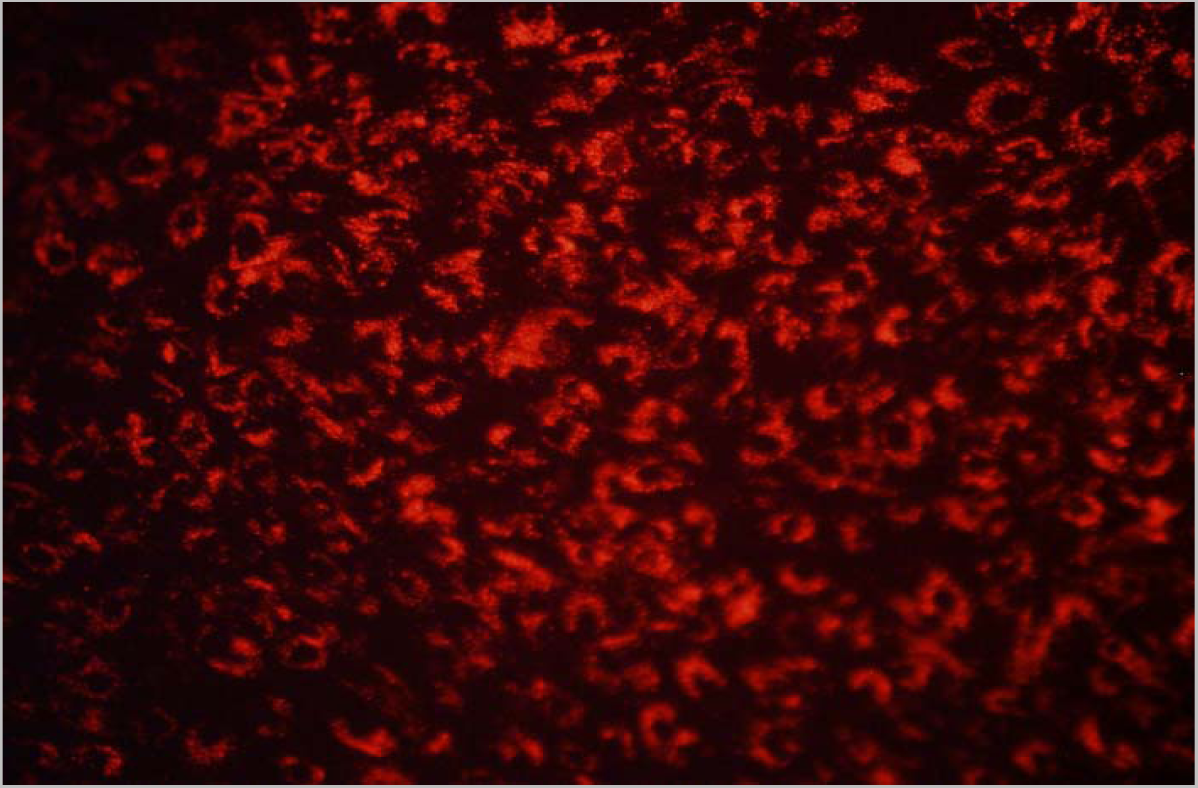
Olfactory cells stained with CM-Dil at the 4th passage (20X)

### 3.4. Formation of multilayer cell pellet

Analysis of the viability and adhesion capabilities of the OTD-SCs from the multilayer cell pellet showed that the resulting cellular mass contained viable cells with high cell adhesion and proliferation properties (Fig. 4).

**Figure 4.**
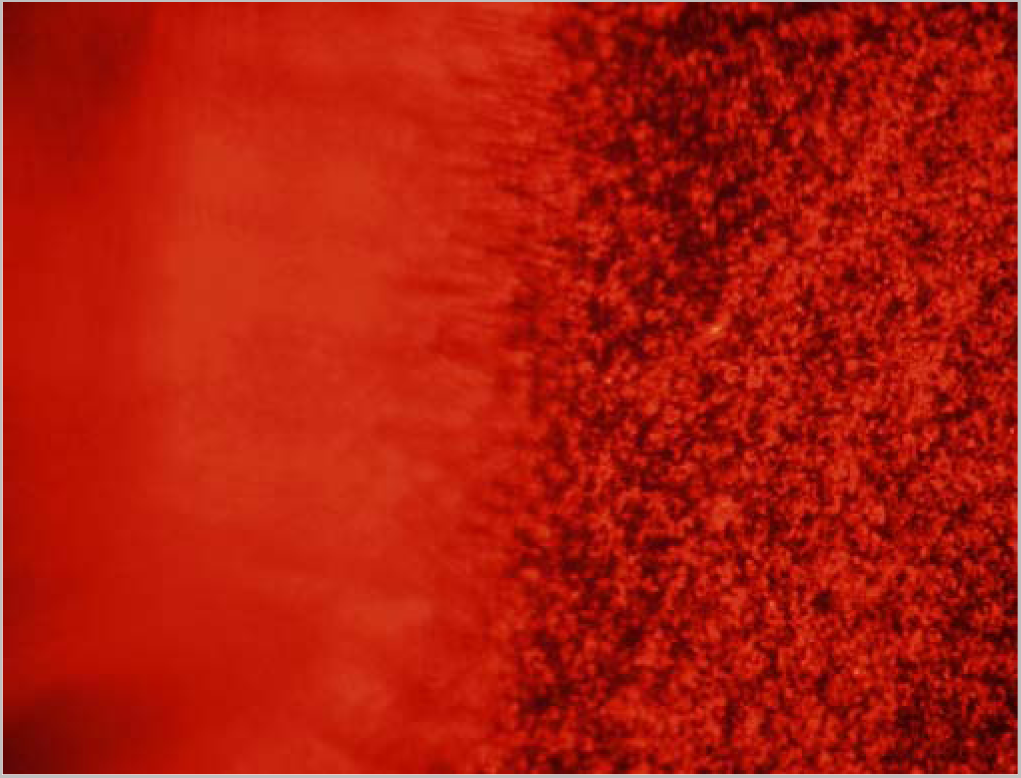
Multilayer cell pellet obtained from olfactory stem cells stained with CM-Dil (10X)

## 4. Discussion

There is a growing research interest in cellular therapies using stem cells for treating difficult injuries that have had unresponsive results with other treatments; one such common and often difficult to treat injury is nerve damage. Previous studies have analyzed the use of stem cells obtained from adipocytes and bone marrow in the treatment of nerve damage (Mantovani et al., 2012; Wang et al., 2012; Liu et al., 2011; Dadon-Nachum et al., 2011; Momin et al., 2010; di Summa et al., 2010; Petrova et al., 2012; Radtke et al., 2011; Wang et al., 2010). However, recent studies show that nerve stem cells obtained from the olfactory region have a higher chance in successful nerve regeneration (Salehi et al., 2009; Shyu et al., 2008). The number of studies investigating these cells, though, is very limited.

In this study, we discuss an efficient isolation of mesenchymal stem cells from olfactory tissue, their culture, characterization, and the formation of a multilayer stem cell mass, to be used in tissue engineering. Biopsies obtained from three male donors of ages 30-45 were used to isolate and culture these cells. The results of this study showed no difference in the cellular and morphological properties of the OTD-SCs obtained from the three different donors. The cells were proven to be olfactory stem cells following morphological, immunophenotypical, and adipocyte differentiation analyses (data not shown). The viability of single and multilayer cell masses was examined both morphologically and by CM-DiI staining. Moreover, the viability of the multilayer cell mass was shown to be preserved for prolonged culture.

In tissue engineering, most stem cell applications require the use of supporting materials. If supporting material is not used, the localization of cells is limited and the resulting nerve regeneration is restricted. The main advantage of using a natural supporting material is that it eliminates the need for expensive and toxic scaffolding materials. A multilayer cell mass can play a crucial role as an efficient and necessary replacement for a supporting material. Due to the interaction of the cells within the multilayer cell culture, the presence of signaling molecules secreted by the cells in their microenvironment and the high number of metabolites supporting cell proliferation are suggested to be more effective in regeneration of damaged nerve cells. No previous studies have been done on this subject. This study showed that forming multilayers maintained the viability and adhesion capacity of the cells rather than having inhibitory effects on cell proliferation. This is the first study to obtain OTD-SCs from a multilayer cell culture. These OTD-SCs are proposed to be used in the treatment of nerve injuries. It is also suggested that metabolites among the multilayer cells allow for better localization in the transplantation. Another advantage of this mesenchymal stem cell mass is that it is cost effective, autologous, and easy to prepare under the appropriate laboratory conditions. These results suggest that placing the multilayer cell mass may allow nerve regeneration even in small areas of damage with no risk of diffusing to other regions.

## 5. Conclusion

The multilayer cells obtained for the regeneration of damaged nerve cells offer better localization in transplantation of the cells than other available methods.

## Acknowledgments

The authors kindly thank Doç. Dr. Özgür Yiğit (Chief Physician at Istanbul Educational and Research Hospital, Kasap İlyas Mah. Org. Abdurrahman Nafiz Gürman Cd. PK: 34098 Fatih / Istanbul) supported for this study. The authors gratefully acknowledge, The Scientific Research Council of Yildiz Technical University (BAP, project number: 2012-07-04 KAP02) for financial support

Original submission.

### Author Disclosure Statement

The authors state that they do not have to disclose any actual or potential conflict of interest including any financial, personal or other relationships with other people or organizations within three years of beginning the submitted work that could inappropriately influence, or be perceived to influence, their work.

